# A closed-loop hand prosthesis with simultaneous intraneural tactile and position feedback

**DOI:** 10.1101/262741

**Authors:** Edoardo D’Anna, Giacomo Valle, Alberto Mazzoni, Ivo Strauss, Francesco Iberite, Jérémy Patton, Francesco Petrini, Stanisa Raspopovic, Giuseppe Granata, Riccardo Di lorio, Marco Controzzi, Christian Cipriani, Thomas Stieglitz, Paolo M. Rossini, Silvestro Micera

## Abstract

Current myoelectric prostheses allow upper-limb amputees to regain voluntary motor control of their artificial limb by exploiting residual muscle function in the forearm^1^. However, the over-reliance on visual cues resulting from a lack of sensory feedback is a common complaint^2,3^. Recently, several groups have provided tactile feedback in upper-limb amputees by using implanted electrodes^4,5,6,7,8^, surface nerve stimulation^9,10^ or sensory substitution^11,12^. These approaches have led to improved function and prosthesis embodiment^4,5,6,7,13,14^. Nevertheless, the provided information remains limited to a subset of the rich sensory cues available to healthy individuals. More specifically, proprioception, the sense of limb position and movement, is predominantly absent from current systems. Here we show that sensory substitution based on intraneural stimulation can deliver position feedback in real-time and in conjunction with somatotopic tactile feedback. This approach allowed two trans-radial amputees to regain high and close-to-natural remapped proprioceptive acuity, with a median joint angle reproduction accuracy of 9.1° and a median threshold to detection of passive movements of 9.5°, which was compatible with results obtained in healthy subjects^15,16,17^. The simultaneous delivery of position information and somatotopic tactile feedback allowed both amputees to discriminate object size and compliance with high levels of accuracy (75.5%). These results demonstrate that touch information delivered via somatotopic neural stimulation and position information delivered via sensory substitution can be exploited simultaneously and efficiently by trans-radial amputees. This study paves the way towards more sophisticated bidirectional bionic limbs conveying rich, multimodal sensations.

---

Despite recent advances in peripheral neuromodulation, direct elicitation of selective proprioceptive percepts remains elusive and is only rarely reported^4,5,7,8^. Efforts to restore proprioceptive feedback invasively have been limited to preliminary studies, showing only modest functional benefits or lacking extensive characterization^18,19,20,21^. Proprioception is thought to be mediated in part by Ia and type II sensory afferents from the muscle spindles^22^. The proximity of proprioceptive afferents and motor neurons within the nerve may explain the difficulty in activating proprioceptive pathways without inducing undesirable motor twitches. Indeed, neurophysiological evidence indicates that microstimulation of proprioceptive afferents does not lead to perceptual responses, unless accompanied by muscle activity^23^. This suggests that selective homologous proprioceptive feedback (i.e., where the restored sensation closely matches the natural sensation, and where there is no co-activation of muscles) could be difficult to achieve with current neural stimulation approaches (in trans-radial amputees). Instead, sensory substitution (remapping) may be a viable alternative, potentially enabling significant functional gains. Sensory substitution has been used extensively in other applications, pioneered by Bach-Y-Rita and colleagues^11,24^, including recently using brain implants in non-human primates^25,26^, and augmented reality in healthy subjects^27,28^, with promising results.

For this reason, we implemented a “hybrid” approach to restore multimodal sensory information to trans-radial amputees, where position information (proprioception) was provided using sensory substitution based on peripheral intraneural stimulation, while pressure information (touch) was restored using a somatotopic approach, where the elicited sensation was correctly perceived on the fingers and palm, as previously shown^4,5^. Specifically, joint angle information was delivered through spared neural afferent pathways using intraneural stimulation of the peripheral nerves in the amputee’s stump. Two transradial amputees were implanted with transverse intrafascicular multichannel electrodes (TIMEs) in the ulnar and median nerves^29^ (Fig. 1). Subject 1 performed a pilot study, while Subject 2 performed a more comprehensive set of experiments. Both subjects reported stable sensations of vibration, pressure, and electricity over the phantom hand and stump during intraneural stimulation (see Extended Data Fig. 1). Position information was provided using active sites which elicited sensations referred to the lower palm area or the stump. This choice avoided any conflict with tactile feedback, which used active sites providing sensations referred to the phantom fingers^4^. The feedback variable was the hand aperture (either one or two degrees of freedom depending on the experiment, see methods), encoded using linear amplitude modulation.

**Figure 1.**
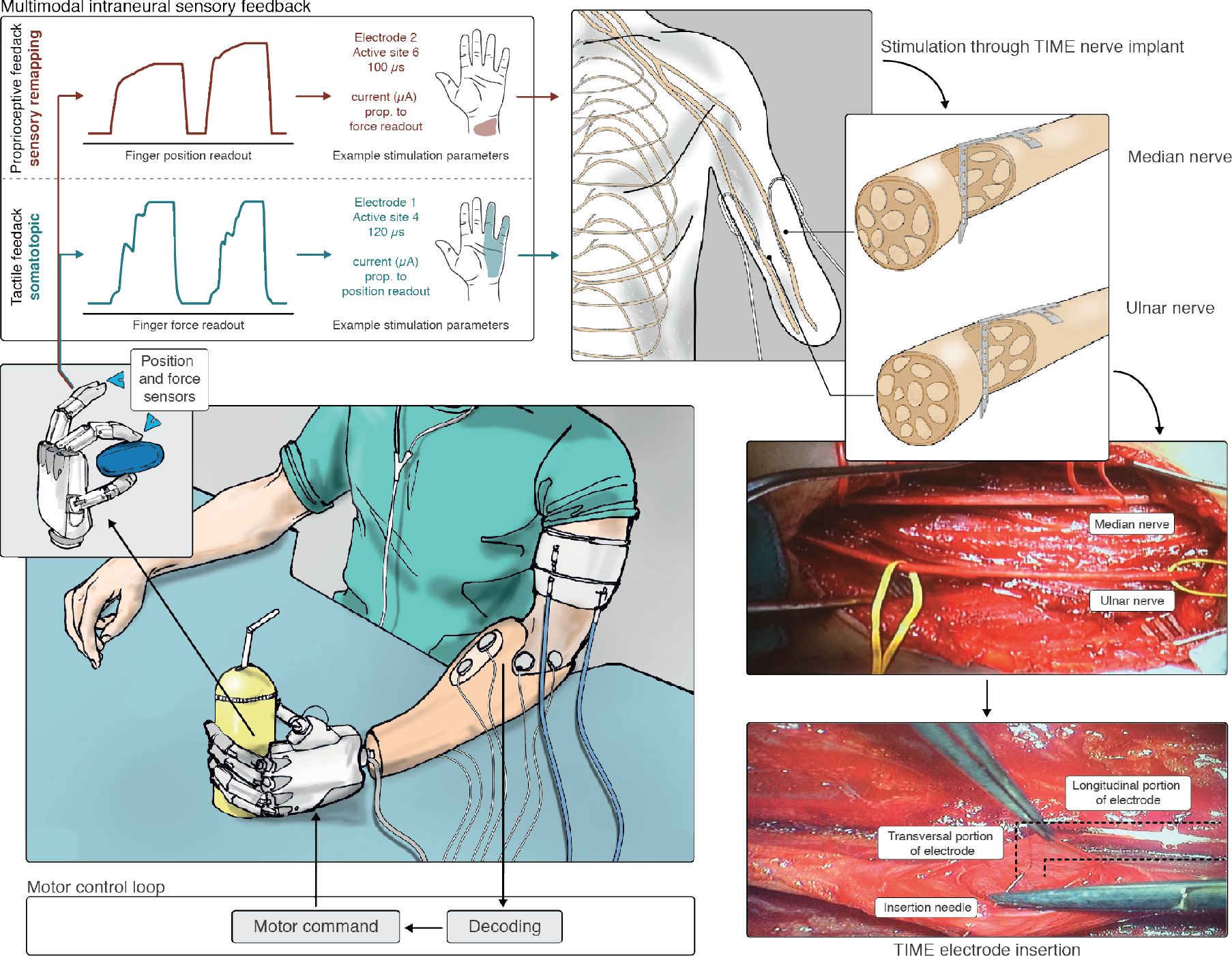
Overview of the multimodal sensory feedback experimental setup. The robotic hand is driven using sEMG activity acquired from the subject’s forearm muscles, and classified into distinct motor commands (bottom left). As the robotic hand closes its fingers around an object, both pressure and position are measured in real-time (bottom left). Information about pressure and position is then encoded into stimulation pulses, where stimulation amplitude is directly proportional to finger position or pressure (top left). Pressure perception is restored using a somatotopic approach, where the induced sensation corresponds to the fingers being touched. Position information (proprioception) is restored using sensory substitution, whereas the sensation does not correspond to the natural area (top left). Both sensory streams are delivered using intraneural stimulation through TIME electrodes implanted in the median and ulnar nerves (top right). The TIME implant is inserted transversally through the exposed nerve fascicles (bottom right).

We first characterized the acuity of the remapped proprioceptive sense alone. We administered two clinical tests, namely threshold to detection of passive motion (TDPM) and joint angle reproduction (JAR)^30^. During the TDPM test, we measured the smallest prosthesis displacement necessary for the subjects to detect passive motion of the artificial hand, starting from randomly chosen positions across the hand’s range of motion. This test measured the sensibility to stimulation amplitude, and is reported in terms of remapped hand aperture. The overall TDPM was 9.5 degrees (interquartile range, IQR = 9.1), with 12.5 degrees (IQR = 10.4) for Subject 1 and 6.5 degrees (IQR = 6.6) for Subject 2 (Extended Data Fig. 2). No statistically significant correlation was found between TDPM and initial hand position (*p* = 0.52), or with movement direction (*p* = 0.11), indicating that proprioceptive sensibility was equal across the range of motion and independent of the direction of movement of the hand (Fig. 2a). Previous results show that healthy individuals obtain TDPM values for single finger joints between 6.5 and 1.5 degrees^15^. Although these results are not directly comparable, it is interesting to note that our approach enabled Subject 2 to obtain an acuity within this range, while Subject 1 obtained a lower acuity. This indicates that the “resolution” of the remapped position sense, determined by the ability to discriminate current amplitudes, may be sufficient for a wide range of functional tasks.

**Figure 2.**
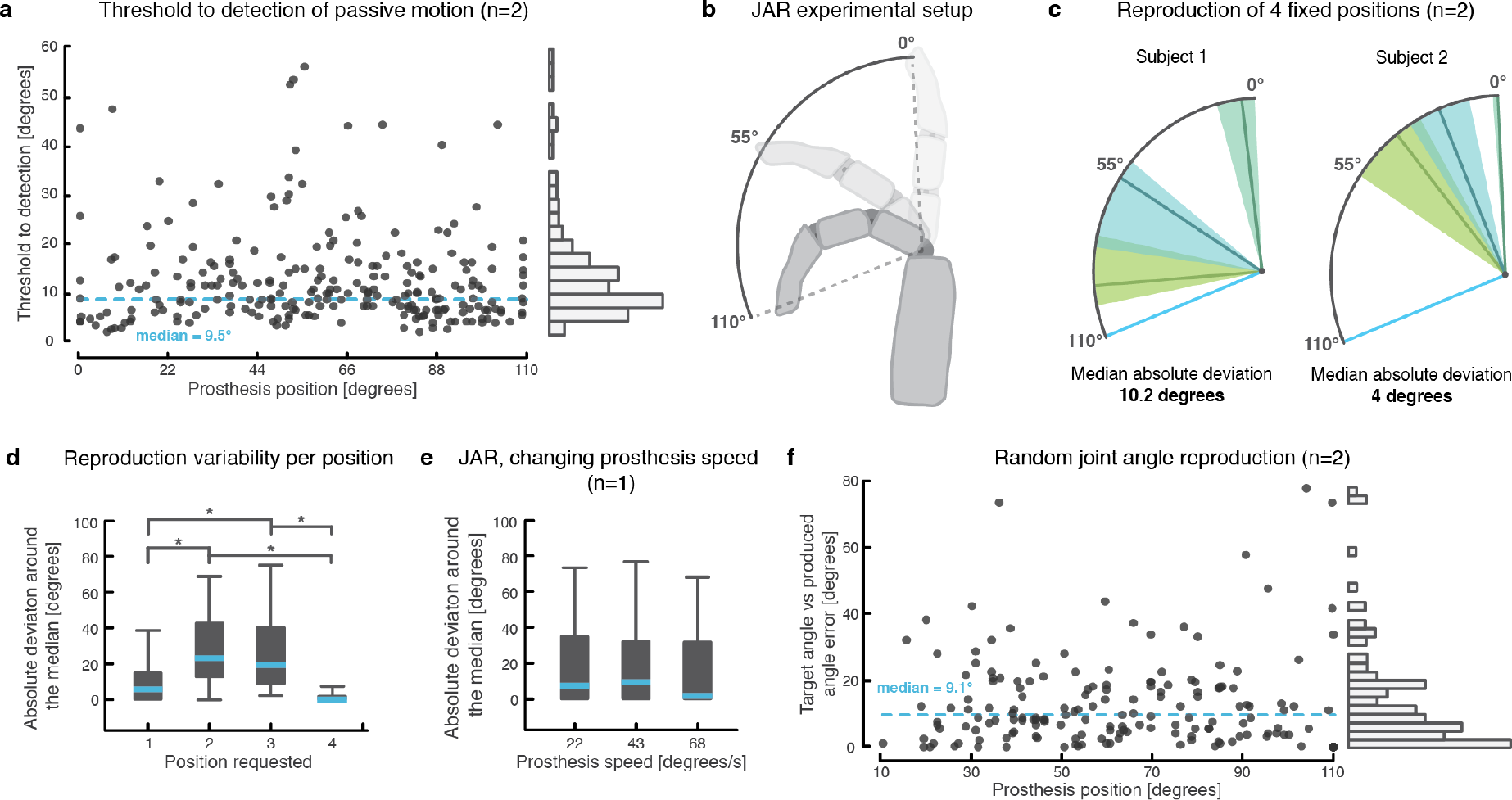
Threshold to detection of passive motion and joint angle reproduction tasks. (**a**) the threshold to detection of passive motion is reported for each prosthesis position tested. The median is reported as a dashed line. A histogram of the data, with bin sizes = 3^°^, is shown on the right. A total of 244 measures were collected with two subjects (115 for Subject 1 and 129 for Subject 2). (**b**) the robotic fingers’ range of motion, and the way the angle is reported. (**c**) JAR accuracy during the fixed position reproduction task for 4 target positions. The reproduced positions are reported as median (full, colored line) and inter-quartile range (shaded area). The median absolute deviation for the pooled performance on all positions is reported for each subject. (**d**) box plots reporting the detailed absolute deviation for each requested position. The median is reported as a blue line, while the box represents the inter-quartile range. The whiskers encompass all data samples (no outliers removed). A total of 80 (40 for Subject 1 and 40 for Subject 2) repetitions were collected for the task. Asterisks indicate conditions found to be statistically different after a Kruskal-Wallis test with multi group correction (**e**) a box plot showing the absolute deviation around the median for randomly switched prosthesis actuation speeds (3 speeds). For this task, only Subject 2 participated, and a total of 48 repetitions were performed. (**f**) a scatter plot shows the measured error in joint angle reproduction for each position tested during the joint angle reproduction task with random and continuous positions. A histogram of the data, with bin sizes = 3^°^, is shown on the right-hand size. A total of 171 measures were collected with two subjects (81 for Subject 1 and 90 for Subject 2).

During a first variant of the JAR test (fixed positions), both subjects were asked to actively move the hand to one of four self-selected angular positions. The angle of closure was measured from the fully open state (Fig. 2b). For each reproduced position, the median absolute deviation from the median (MAD, a robust measure of variability) was computed. MAD was measured at 10.2 degrees for Subject 1, and 4 degrees for Subject 2 (Fig. 2c). Overall, MAD was significantly lower when the target position was at the extremes of the range of motion (fully open or fully close) compared to intermediate positions due to the impossibility to “overshoot” the target at both extremes of movement (*p*<0.05, Fig. 2d).

Subject 2 also performed a control condition to dismiss the possibility of using movement duration to infer finger position. Indeed, during the same task, hand prosthesis actuation speed was randomly switched between three values (22, 43, and 68 degrees/s), without the subject’s knowledge. Despite receiving unreliable information about timing, no significant increase in spread was observed for any of the tested actuation speeds (*p* = 0.76), nor for the overall performance (*p* = 0.75), indicating that timing did not play a critical role in achieving high task performance (Fig. 2e).

We also performed a more challenging JAR experiment using random and continuous positions. In this case, the robotic hand was first passively closed with a random joint angle. Then, the hand was passively opened again, and the subjects were asked to control the robotic hand and bring it back to the same position. The JAR accuracy was constant across the entire range of motion (*p* = 0.68), with a median error of 9.1 degrees (IQR = 14.6) (Fig. 2f). Median error was 8.6 degrees for Subject 1 (IQR = 12.7) and 9.9 degrees for Subject 2 (IQR = 15.9) (Extended Data Fig. 2). Despite the imprecision introduced by the controller delay (approximately 100ms), these errors compare favourably to results obtained with healthy individuals (matching error for the metacarpophalangeal joint was measured between 5.94 degrees and 10.9 degrees for healthy subjects^16,17^).

To study how the remapped position sense could be exploited during functional tasks, we performed an object size identification experiment, where subjects had to determine the size of an object chosen randomly from a pool of four cylinders with varying diameter (Fig. 3a). The objects resulted in different final degrees of closure of the hand (Fig. 3d). Overall, the two subjects identified the objects correctly in 78% of cases (77.5% for Subject 1 and 80% for Subject 2, Extended Data Fig. 3), while five healthy controls had a higher score of 98.5% (Fig. 3b, and Extended Data Fig. 4a). Supplementary video S1 shows a few example trials of the object recognition task.

**Figure 3.**
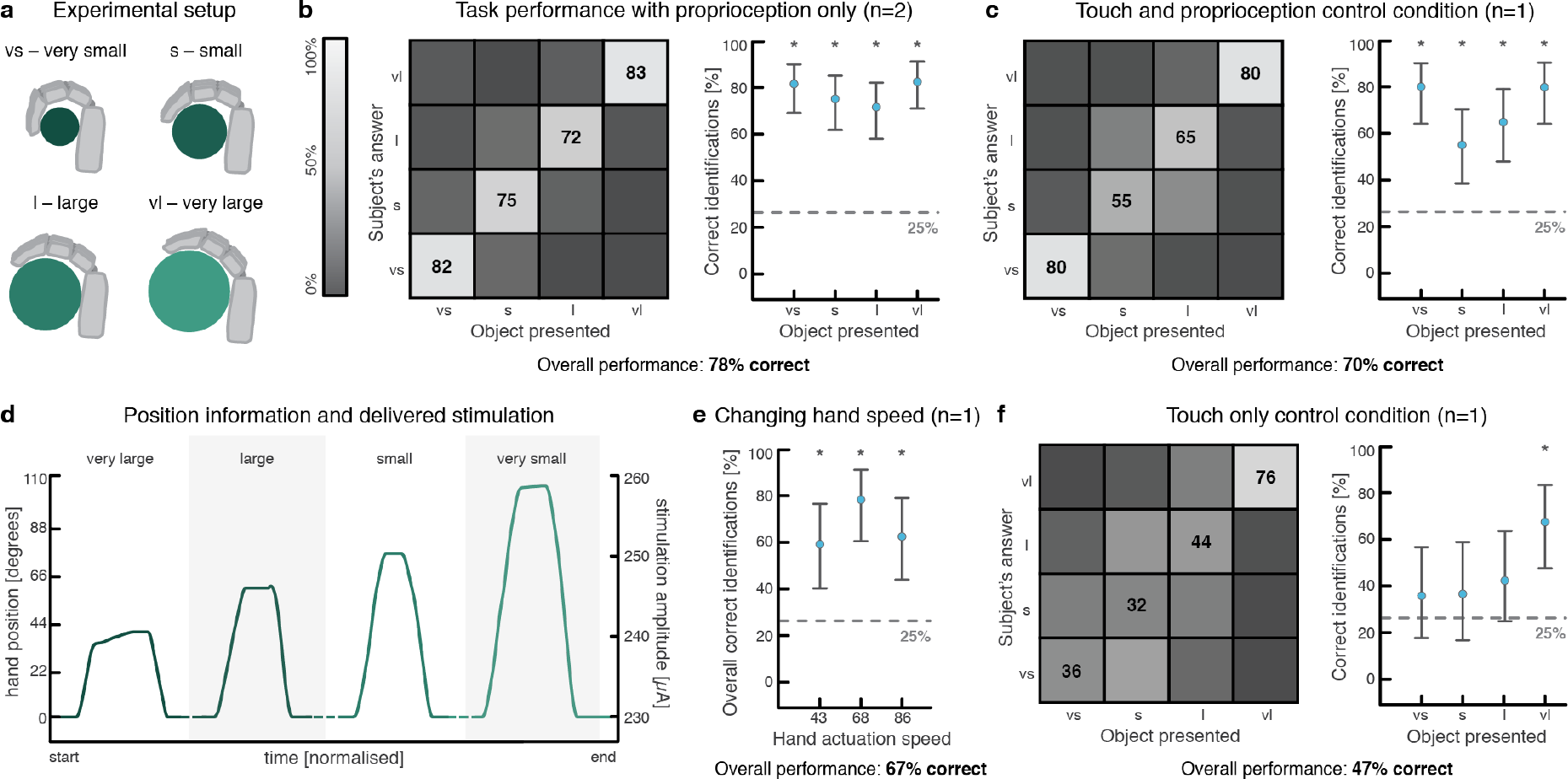
Identification of object size. (**a**) schematic representation of the four different objects used during the object size identification task, and their labelling (not to scale). (**b**) overall performance during the task with remapped proprioception only for the amputee subject in the form of a confusion matrix (left) and performance in identifying each object (right). Median correct identifications and a 95% confidence interval for each object are reported alongside the matrix. Stars identify levels which were statistically different from chance level. A total of 160 repetitions (40 for Subject 1 and 120 for Subject 2) were performed with two amputee subjects. (**c**) overall performance during the object size recognition task with simultaneous touch and proprioceptive feedback in the form of a confusion matrix. A total of 100 repetitions were performed with Subject 2. (**d**) representative position traces obtained during the experiments. One example was chosen for each cylinder size, to illustrate the difference in measured position obtained in each case. In addition, the stimulation amplitude computed from the position is reported on the second y-axis. (**e**) overall performance for each tested hand actuation speed during a control trial with changing speeds. A total of 96 repetitions were performed with Subject 2. (**f**) the performance obtained during a control condition where only touch feedback was delivered. In this case, 100 repetitions were performed with Subject 2.

Several control conditions were tested with Subject 2. First, the same task was repeated with tactile feedback alone (Fig. 3f). In this scenario, performance was poor, but remained above the 25% chance level (47% correct identification, 95% CI [36.9, 57.2]). However, further analysis showed that only the largest object was correctly identified above chance level (Fig. 3f). Second, the task was performed with both tactile and remapped position feedback. The measured performance (70% correct identification) was not statistically lower than the performance obtained with remapped proprioception only (*p* = 0.449, Fisher’s exact test), indicating that the addition of touch did not interfere with the interpretation of position feedback (Fig. 3c). Third, when remapped proprioception was provided alone, and the prosthesis movement speed was randomly switched between three values, the performance was 67%, which was not statistically different from the condition with constant speed (*p* = 0.226, Fisher’s exact test, Fig. 3e).

Data obtained with Subject 2 for the object size task shows a steady increase in performance over time, indicating that although remapped proprioception can successfully be exploited almost immediately, training may confer an advantage, and could lead to further improvements in performance over time (Extended Data Fig. 5a). Additional measurements, obtained over longer periods of time, could confirm the effect of training on performance.

Both subjects were also asked to identify the size and compliance of four different cylinders (Fig. 4a). In this case, tactile and proprioceptive feedback was provided simultaneously (Fig. 4d). Overall, performance for this task was high, with 75.5% correct answers (87.5% for Subject 1 and 73% for Subject 2) (Fig. 4b and Extended Data Fig. 3). By comparison, two healthy controls had a perfect score of 100% (Extended Data Fig. 4b). Subject 2 performed the same task while receiving only remapped position (Fig. 4c) or tactile (Fig. 4e) feedback. In both cases, performance significantly worsened (no overlap of 95% confidence intervals). Interestingly, when only position feedback was provided, object size was identified above chance level, while object compliance was not (Fig. 4c). Conversely, when tactile feedback was provided, only object compliance was correctly identified (Fig. 4e). This shows that each sensory modality mainly provides information regarding one object feature (touch informs about compliance^4^, and position feedback about size). Furthermore, providing both modalities simultaneously can improve performance, as seen from the superior compliance decoding achieved using both touch and proprioception compared to either modality individually, for the large object (Fig. 4f).

**Figure 4.**
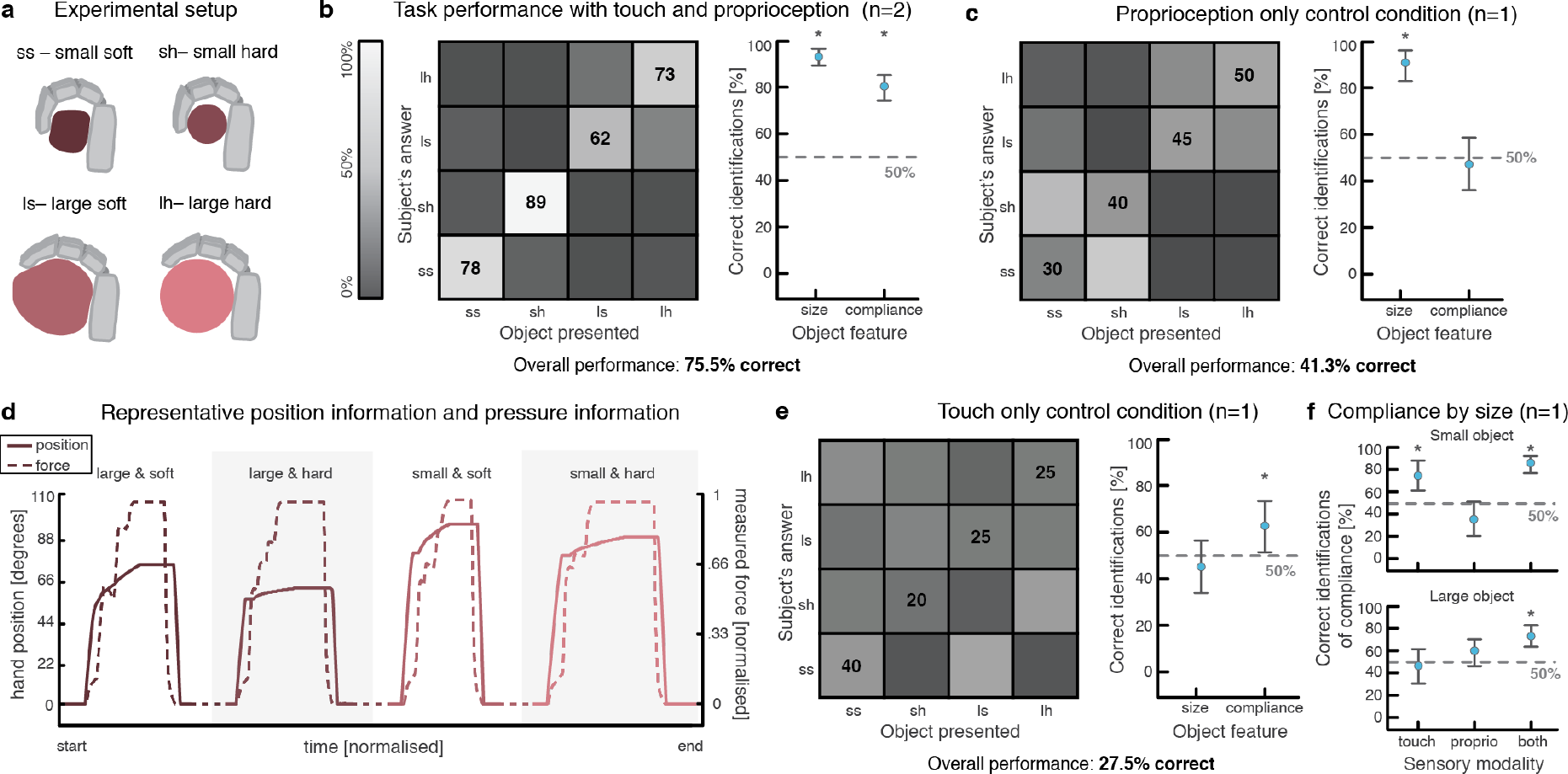
Identification of object size and compliance. (**a**) schematic representation of the four different objects used during the object size and compliance task, and how they were labelled (not to scale). (**b**) overall task performance with both remapped proprioception and touch, for both subjects, reported as a confusion matrix. The combined performance is shown under the image. Median correct identifications and a 95% confidence interval for each object feature (size and stiffness) are reported alongside the matrix. Stars identify levels which were statistically different from chance level. A total of 220 repetitions were performed with two subjects (40 for Subject 1 and 180 for Subject 2). (**c**) performance during the same object size and compliance task when only proprioceptive feedback is provided. A total of 80 repetitions were performed with Subject 2. (**d**) representative force and position traces, as measured by the robotic hand, for each object type. The full lines represent hand position (0°-110°), while the dashed lines represent measured force (normalized). The four patterns are not contiguous (illustrated by dashed lines), but the relative duration of each pattern is conserved, to allow meaningful comparison of the slopes. (**e**) performance during the same task when only touch feedback is provided. A total of 80 repetitions were performed with Subject 2. (**f**) compliance decoding performance broken down by object, with touch only, proprioception only, or both sensory modalities. Compliance decoding performances above chance level are shown with a star. A total of 380 repetitions were used for this panel (combination of data from **b**, **c** and **e**).

In another experiment, we provided two channels of remapped proprioceptive feedback simultaneously (one channel for the first three digits and one for the last two). In this case, two channels giving rise to different sensations on the stump were used. Using this “multijoint” feedback, Subject 2 could simultaneously detect the diameter of two cylinders with a very high performance of 93.7% (Extended Data Fig. 6), demonstrating that the sensory remapping approach presented here can also be applied to more than one finger simultaneously.

This study shows that trans-radial amputees can effectively exploit a hybrid multimodal stimulation approach, which combines somatotopic feedback (i.e., touch) with sensory substitution (i.e., remapped proprioception). The functional results demonstrate that the two streams of information can be used simultaneously to achieve high task performance. Our results pave the way towards more sophisticated bidirectional bionic limbs conveying rich, multimodal sensations.

**Supplementary information** is available alongside this paper.

The data that support the findings of this study are available from the corresponding authors upon reasonable request.

The Matlab code used for the analysis of the results presented in this study is not available as it was scripted in an interactive session, and can readily be replicated by other researchers. The custom C++ code used to control the various hardware components is not made available, as it is too specific to the exact set of hardware equipment (e.g. brand, model, version), as described in the methods section. It is our opinion that this software would not be useful to other researchers in its current state.

## Author contributions

E.D., G.V., A.M., and S.M. conceived the experiments. E.D., G.V., I.S., A.M. and J.P. conducted the experiments, E.D. analyzed the results, E.D. created the figures, E.D. and S.M. wrote the manuscript. E.D., G.V., I.S., S.R. and F.P. characterized the sensations elicited by stimulation during the experiments. G.G., R.I. and P.M.R. recruited the patient and were responsible for all the clinical activities. M.C. and C.C. developed the sensorized hand prosthesis, and T.S. developed the TIMEs. All authors reviewed the manuscript.

## Methods

### Patient recruitment and experiment logistics

Two amputees participated in the study (a 54-year-old female with a left wrist disarticulation incurred 23 years prior to the study, and a 54-year-old female with a proximal left trans-radial amputation incurred 2 years prior to the study). Ethical approval was obtained by the Institutional Ethics Committees of Policlinic A. Gemelli at the Catholic University, where the surgery was performed. The protocol was also approved by the Italian Ministry of Health. Informed consent was signed. During the entire length of our study, all experiments were conducted in accordance with relevant guidelines and regulations. This study was performed within a larger set of experimental protocols aiming at the treatment of phantom limb pain and robotic hand control. The clinical trial’s registration number on the online platform www.clinicaltrials.gov is NCT02848846.

The data reported in this manuscript was obtained over a period of several days in two amputees. The first patient (Subject 1), was recruited as a pilot case towards the end of an ongoing long-term study of intraneural electrodes (5 months after implantation), and performed a more limited number of experiments (particularly with regards to control conditions). Subject 1 performed all experiments reported here over a period of four days (divided in two sessions of 2 back-to-back days over two weeks), although each type of experiment was not performed more than once (there is no data for the same experiment over multiple days). The second patient, (Subject 2), was recruited at an earlier stage (2 weeks after implantation), performed a larger number of trials and a more complete set of control experiments. All data for Subject 2 was obtained over a period of 6 days (sessions spread over a period of six weeks), with several experiments grouping data over multiple days (and allowing a comparison of performance over days, as shown in Extended Data Fig. 5a).

### Bidirectional setup and prosthesis control

For the functional tasks, subjects were fitted with a custom bidirectional research prosthesis, allowing control of hand opening and closing by processing surface electromyographic (sEMG) signals, and providing sensory feedback by means of electrical stimulation of the peripheral nerves. A robotic hand with tension force sensors integrated within each digit (IH2 Azzurra, Prensilia, Italy) was controlled using a custom, multithreaded C++ software running on a RaspberryPi 3 single board computer (Raspberry Pi Foundation, UK). A recording and stimulating device (Neural Interface Processor, Ripple, LLC, US) was also connected to the central single board computer, acquiring sEMG data from two or four bipolar channels, and providing stimulation outputs to the four neural electrodes. Custom moulded sockets were built with integrated screws to easily fix the robotic hand on the end. Holes were drilled to allow for the placement of sEMG electrodes on the stump.

For prosthesis control, a simple 3 state (open, close, rest) threshold controller was used for Subject 1, and Subject 2 used a KNN (k=3) classifier with 3 classes^31^. Two or four bipolar channels of sEMG were acquired from forearm residual muscles (for Subject 1 and 2 respectively), where palpation was used to place the electrodes in the optimal positions. The sEMG data were acquired with a sampling frequency of 1 kHz, and filtered using an IIR filter with 4^th^ order Butterworth characteristics, between 15 and 375 Hz, as well as a notch filter to remove 50 Hz power hum. For the threshold controller, the mean absolute value (MAV) was computed for each channel, and a threshold was set manually to indicate when the hand should be opened or closed. the amplitude of the sEMG signal (MAV) controlled hand actuation speed (proportional control). For the KNN classifier, the waveform length was computed over a window of 100ms for each channel and fed to the classifier every 100ms. The decoded class was used to send open or close commands to the prosthesis.

### Tactile feedback based on intraneural electrical stimulation

Both subjects were implanted with four TIMEs in the median and ulnar nerves (two per nerve), above the elbow, each with 14 active sites and two counter electrodes on the substrate^29^. A total of 56 actives sites per subject were thus available. After an extensive mapping phase, during which the stimulation parameter space (defined by the following variables: electrode, active site, stimulation amplitude, stimulation pulse width and frequency) was explored, a relationship between stimulation parameters and sensation quality, location and intensity was established, as described in Raspopovic et al., where an analogous preparation was used^4^. Briefly, for every active site, injected charge is increased progressively at a fixed frequency and pulse width, by changing the stimulation amplitude. If the range afforded by the selected pulse width and the maximum deliverable current amplitude (imposed by the stimulator) is too small, the pulse width is incremented and the same thing is done again. The threshold for minimum sensation is noted as soon as the subject detects any sensation related to the stimulation. The maximum parameters are saved when the sensation becomes painful, starts inducing a muscle twitch or simply if the patient does not feel comfortable increasing it further. This is repeated three times per active site, giving an average value. These two values, threshold and maximum, are saved for every active site, and can later be used when choosing a modulation range. The effects of changing frequency were not investigated in this work, and it was always fixed at 50Hz. Injected current levels were always below the chemical safe limit of 120nC for each stimulation site.

During the experiments reported in this work, a single tactile channel was used for sensory feedback in both subjects at any given moment (the optimal electrode and active site for the experiments were chosen every week based on the sensations reported by the subjects, and were not always the same). The measured force applied by the prosthetic digits was encoded using a linear amplitude modulation scheme, designed to associate perceived stimulation intensity with measured force. Parameters were chosen in such a way as to optimally cover the whole dynamic range of sensations reported by each subject. For Subject 1, tactile feedback was provided using charge-balanced, square pulses with an amplitude between 230μA and 500μA, and a pulse width duration between 80μs and 120μs, which resulted in a sensation of vibration referred to the base of the middle finger. For Subject 2, tactile feedback was provided using an amplitude between 90μA and 980μA, and a pulse width duration between 50μs or 200μs depending on the day, which always resulted in a sensation of pressure or contraction referred to most of the ulnar innervation area (although less intense over the fourth finger). A more detailed set of parameters is provided in the Extended Data Fig. 1b. Only two sets of parameters were used simultaneously at any given time (one for each feedback channel). The high number of parameters reported in the table are a result of changes in parameters between days and sessions, especially for Subject 2 who performed these experiments soon after implantation, when stimulation parameters may still vary significantly from day to day. Indeed, the mapping procedure was repeated every week, often leading to the discovery of better active sites and sensations which were then used during the experiments.

For the typical time scales involved in our experiments (trials lasting in the order of minutes), neither of our subjects reported relevant changes in sensation intensity, which would indicate the presence of adaptation. Indeed, such effects were anecdotally observed only for much higher stimulation durations (tens of minutes). For all practical purposes, adaptation was insignificant during our experiments.

### Sensory substitution for proprioceptive feedback

To convey position information to the subjects, sensory substitution was employed. To avoid any cross-talk with tactile feedback, an active site resulting in a sensation which was not referred to the fingers was used. In Subject 1, the selected stimulation parameters resulted in paraesthesia located in the lower palm area, while in Subject 2, the area involved was the medial part of the forearm, occasionally extending into the lower palm and wrist. Encoding of the position information retrieved from the robotic hand (a value between 0-255, corresponding to hand aperture angles of 0-110 degrees, as measured on the robotic hand) was achieved using a simple linear encoding scheme. After establishing a suitable modulation range for each selected active site, the hand position value was used to modulate stimulation amplitude, while pulse-width and frequency were kept constant (f = 50Hz). Amplitude modulation resulted in changes to the perceived sensation intensity. The range of parameters used for stimulation were as follows: an amplitude between 230μA and 260μA, and a pulse width duration of 80μs for Subject 1, and an amplitude between 100μA and 600μA, and a pulse width duration of 100μs or 200μs depending on the day for Subject 2.

Both subjects underwent a brief learning session (<20 min) to help map the stimulation intensity to the prosthesis opening angle. We first instructed each subject to explore the new information by looking at the robotic limb while it was passively opened and closed. Then, we turned the control on and instructed the subjects to actively explore their environment, grasping various objects and performing opening and closing movement with the prosthesis. Both subjects quickly expressed confidence in interpreting the sensation, as well as a readiness to initiate the trials. Over the entire duration of the trials, the subjective experience associated with the remapped proprioceptive stimulation remained constant (perceived as paraesthesia or contraction respectively).

### Threshold to detection of passive motion

During the TDPM task, the robotic hand was moved passively using a software interface controlled directly by the experimenter. Proprioceptive stimulation was provided during the entire trial. The subjects were instructed to announce when a movement was felt, and in what direction. Whenever a movement was detected, the initial position and the detection position were saved. Then, after a small pause, the experiment continued starting from the last position. During these experiments, the subjects were acoustically and visually isolated, using a sleeping mask and a set of headphones playing music. Falsification trials with no stimulation were also carried out. Prosthesis actuation speed was 27.5 deg/s. To eliminate the possibility that time of actuation was being used as a proxy for degree of closure, a random amount of time was used between each repetition. Thus, the time between the beginning of each trial and the first movement of the hand was not fixed. For Subject 2, the control algorithm for passively moving the hand was modified to ensure that each time the experimenter requested a “step” in each direction (open or close), the resulting movement would be of lower magnitude (1.25 degrees, fixed). Indeed, the setup used for Subject 1 was found to be inaccurate (step size of 9.5±5.5 degrees), and this was improved for the second set of experiments. In other words, in the case of Subject 1, when passively moving the hand, the experimenter could not generate movements smaller than 9.5^°^ on average. Consequently, if the “true” TDPM accuracy was lower than this (as it was found to be for Subject 2) our experimental setup would not have been accurate enough to measure it. This is an important limitation to keep in mind when looking at the TDPM results for Subject 1.

### Joint angle reproduction

We performed two variants of the JAR test. In the first variant, the subjects were instructed to bring the hand to one of four self-selected positions. Before starting the experiments, we asked each subject to show us the chosen positions using their intact hand. This was done to ensure they had understood the task. During the rest of the trial, the positions were recalled from memory. For every requested position, the final position of the hand was recorded, and after a brief pause, the next position was requested (the same position was never asked twice in succession). Subject 2 performed an additional set of control trials, where the prosthesis actuation speed was randomly drawn from a set of three possible speeds (22, 43 and 68 degrees/s).

In the second JAR variant, there were no pre-defined positions. Instead, the robotic hand was closed to a random and continuous position passively by the experimenter, and the subjects could “feel” the sensation for a few seconds. The hand was then opened again, and the subjects were instructed to bring it back to the same position actively. Both the initial position and the reproduced position were recorded, before the next repetition would start. During all variants, the subjects were acoustically and visually isolated, as described above. Additionally, falsification trials with no stimulation were carried out. As with the TDPM task, the time between the beginning of each trial and the first movement of the hand was not fixed during the last JAR variant (this was impossible during the first variant, since the trial was initiated by the subject).

### Object size identification

During the size identification task, four 3D printed cylinders of equally spaced diameters were used (2cm, 4.33cm, 6.66cm and 9cm, referred to as sizes very small, small, large, and very large, respectively). The choice of four cylinders was based on pilot results which indicated that using a smaller number would result in the task not being challenging enough (Extended Data Fig. 5b). After being acoustically and visually isolated, both subjects were asked to close the robotic hand, while one of the four objects was placed in its grip. The subjects announced which object was thought to be held in the hand, and both the actual object and reported object were recorded. A simple control trial with no stimulation was also carried out. Additionally, Subject 2 performed a series of control trials. First, the same task was carried out using only tactile feedback. Second, the task was performed with both tactile and proprioceptive feedbacks together. Finally, the task was performed while prosthesis actuation speed was randomly drawn from a set of three possible speeds (43, 68 and 86 degrees/s).

To establish a baseline accuracy of natural hand proprioception, five right-handed healthy subjects were recruited to perform the same size recognition task. Their right arms were placed in a fixed position on a table, allowing for palmar grasps. To more closely match the experiment performed with the robotic limb, the objects were presented in such a way that they would not touch the thumb, being wedged instead between the fingers and the palm.

### Combined size and compliance identification

The combined size and compliance identification task was performed the same way as the object size identification experiment described above. Here, the objects had two different sizes and two different compliances (hard 3D printed plastic and soft foam), allowing a total of four different combinations. In addition to the proprioceptive stimulation provided in all the other experiments, touch feedback was delivered by means of electrical nerve stimulation during this trial. Both subjects performed this task. In addition to the base task, Subject 2 performed two control conditions. In the first, the same task was performed while only proprioceptive feedback was turned on. In the second, the same was done with only tactile feedback turned on.

To establish a baseline accuracy of natural hand proprioception, two right-handed healthy subjects were recruited to perform the same size recognition task.

### Multi-joint proprioception task

Instead of providing one channel of tactile feedback and one channel of proprioceptive feedback, as in previous tasks, Subject 2 also performed a task where two channels of proprioceptive feedback were provided simultaneously. In this case, one channel encoded the degree of closure of the median area (first three fingers), while the second channel encoded the degree of closure of the ulnar area (last two fingers). In this case, two channels giving distinct sensations on the forearm were used, with the same overall approach described above. With this multi-joint feedback, Subject 2 was asked to recognize four conditions: two small objects placed in the median region and ulnar region, a small object placed in the median region and a large object placed in the ulnar region, the opposite condition with the small object in the ulnar region and a final condition with two large objects. The rest of the task’s details were kept identical to the object size experiment described above.

### Statistics and data analysis

All data was analyzed using Matlab (R2016a, The Mathworks, Natick, US). All statistics were performed using the available built-in functions. A one-sample Kolmogorov-Smirnov test was used to determine if the datasets associated with the various experiments were normally distributed. None of our datasets passed the test as they are highly asymmetrical due to the nature of the tasks. We therefore used non-parametric alternatives (Kruskal-Wallis instead of Anova) and reported the median and inter-quartile range instead of the average and standard deviation. All reported *p*-values resulting from Kruskal-Wallis tests measure the significance of the chi-square statistic. When appropriate, multi group correction was applied using Tukey’s Honestly Significant Difference Procedure *(multcompare(),* Matlab). Spearman’s rank correlation coefficient was used to test if the scatter plots shown in Fig. 2 had correlation values significantly different from 0. Spearman’s rank correlation was used instead of Pearson’s linear correlation coefficient which assumes normality. To measure the spread of data in the JAR experiments (Fig. 2), the robust and non-parametric median absolute deviation from the median (MAD) was used. In Fig. 2f, a Kruskal-Wallis statistic was computed to test the hypothesis that the measured deviation was dependent on the position tested. A multiple comparison correction was applied. Levels that were found to be statistically different are marked with a star (*p* < 0.05). All plots representing median and 95% confidence interval in Fig. 3b, c, e and f and Fig. 4b, c, e and f were generated using a binomial parameter estimate, with chance level being estimated at 25% for correctly recognizing one amongst four objects, and 50% for correctly identifying one feature (hard vs soft, big vs small). Non-overlap of 95% confidence intervals was used as a sufficient criterion to identify statistically significant differences, while a Fisher’s exact test was used in cases where it was not possible to draw conclusions directly from the intervals (i.e. 95% confidence interval overlap). Additional details about the number of repetitions for each experiment are reported in the corresponding figure legends. When random numbers were needed (e.g. generating object presentation sequences), random permutations of an equi-populated sequence (*randperm()*, Matlab) were used.

## Extended Data Figure

**Extended Data Figure 1.**
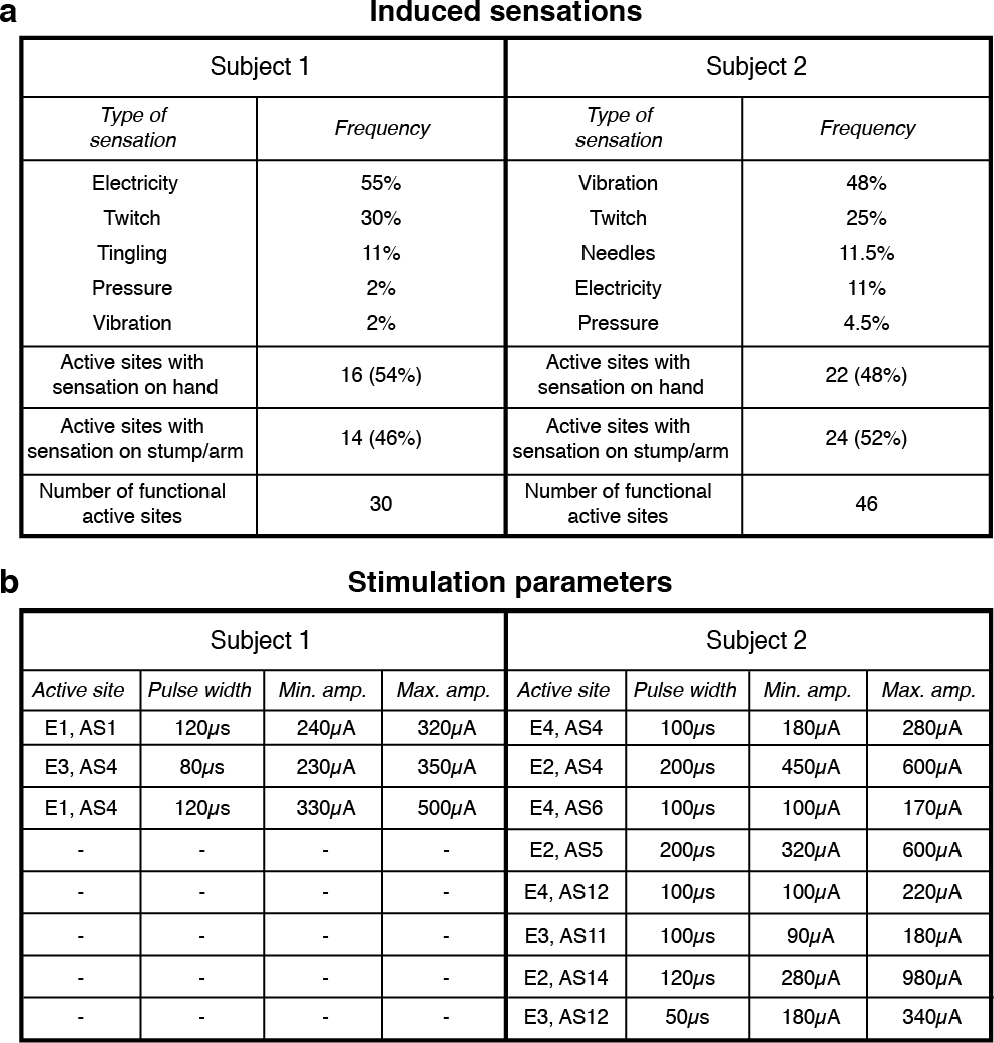
Table of the reported induced sensations and stimulation parameters. (**a**) general information about the intraneural stimulation induced sensations. The occurrence frequency of each type of sensation quality over all active sites is reported, as well as the number of functional active sites and the proportion of active sites giving rise to sensations in the stump and in the phantom hand. (**b**) each set of stimulation parameters used during the experiments (only a subset of all available active site). E refers to the electrode number (out of four) and AS refers to the active site (14 per electrode). The experiments were performed with the same parameters within sessions, but parameters sometimes changed between days, leading to a high number of different combinations used over the entire duration of the experiments.

**Extended Data Figure 2.**
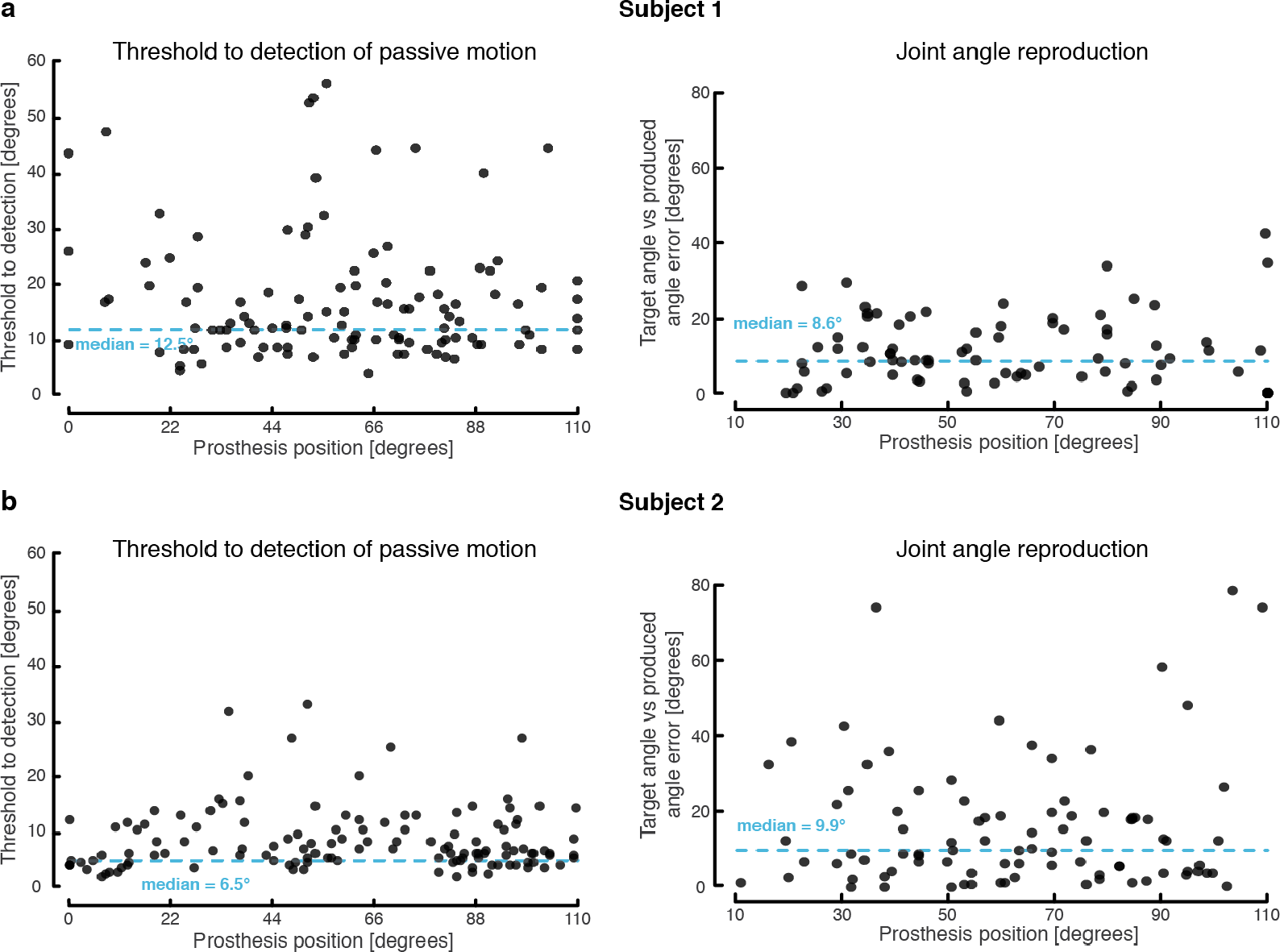
TDPM and JAR performance broken down by subject. (**a**) TDPM (left) and JAR (right) measures for Subject 1, presented in the same format as Figure 2, without the histogram. A total of 115 measures were collected for the TDPM task, and 81 measures were collected for the JAR task. (**b**) TDPM (left) and JAR (right) measures for Subject 2, presented in the same format as Figure 2, without the histogram. A total of 129 measures were collected for the TDPM task, and 90 measures were collected for the JAR task.

**Extended Data Figure 3.**
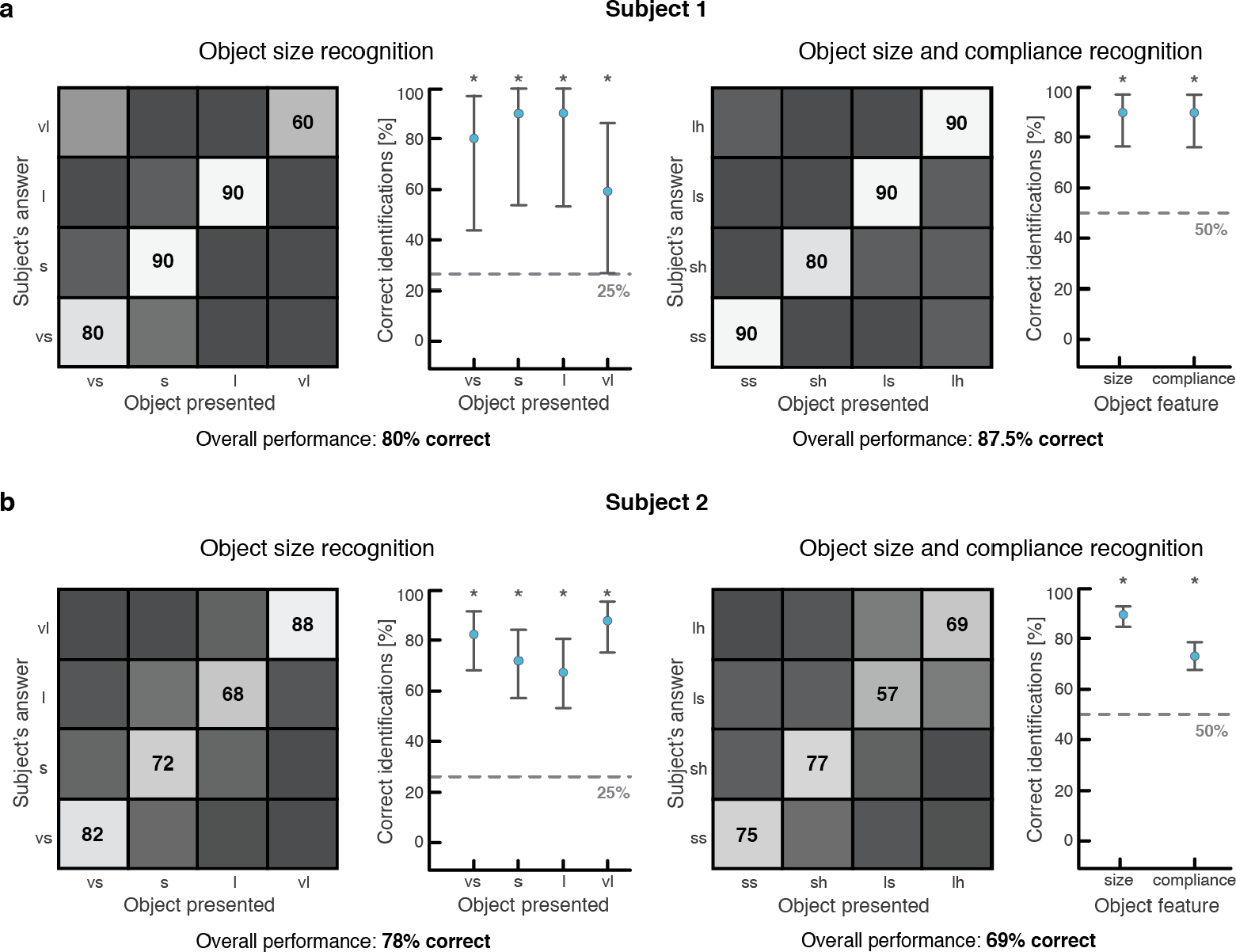
Object size and compliance recognition broken down by subject. (**a**) Object size (left) and object size and compliance (right) tasks performances for Subject 1, presented in the same format as Figure 3 and 4. A total of 40 measures were collected for the object size recognition task, and 40 measures were collected for the object size and compliance recognition task. (**b**) Object size (left) and object size and compliance (right) tasks performances for Subject 2, presented in the same format as Figure 3 and 4. A total of 120 measures were collected for the object size recognition task, and 180 measures were collected for the object size and compliance recognition task.

**Extended Data Figure 4.**
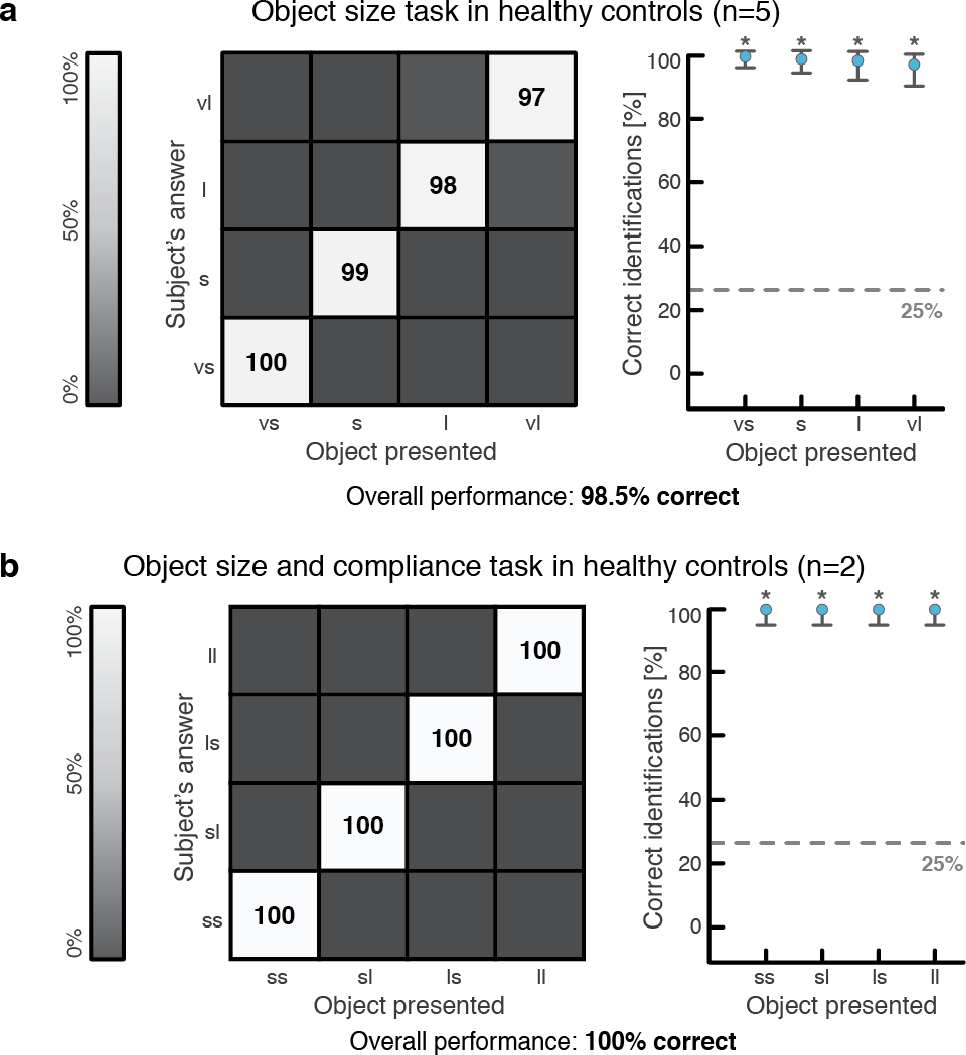
Performance of healthy controls during object identification tasks. (**a**) the object size identification experiment was performed with 5 healthy subjects, with 80 repetitions performed with each healthy control, leading to a total of 400 repetitions. (**b**) the object size and compliance identification task was performed with 2 healthy subjects, with 80 repetitions performed with each healthy control, leading to a total of 160 repetitions.

**Extended Data Figure 5.**
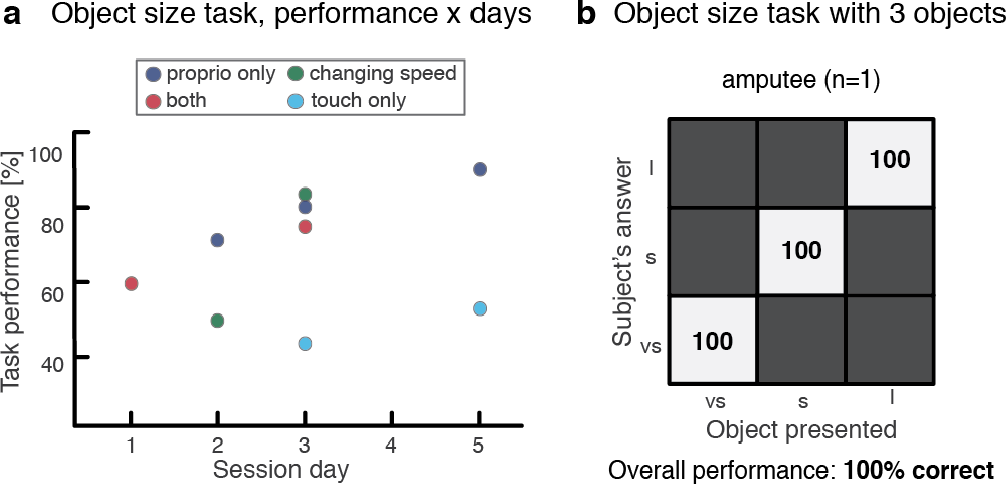
Control condition and time progression of object recognition tasks. (**a**) overall performance over the various days for each type of object recognition task variant (with control conditions). The results reported are for Subject 2. The first subject did not perform the same experiment on multiple days. Only a subset of the experiments was performed each day, as shown. (**b**) confusion matrix reporting the performance of the object recognition task with only three objects, reported for a single subject, with 45 repetitions.

**Extended Data Figure 6.**
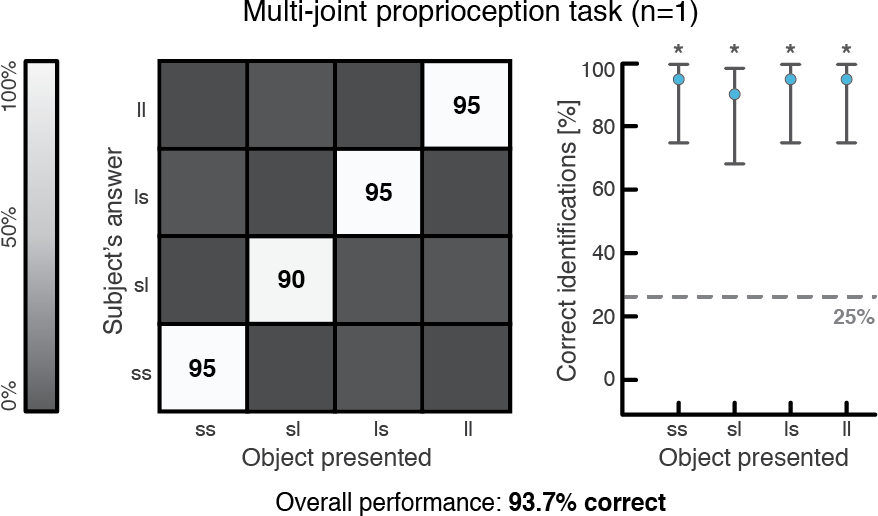
Multi-joint proprioceptive task performance. Confusion matrix showing the overall performance measured during the multi-joint proprioception task. The labels indicate the type of objects presented; ss: small and small, sl: small and large, ls: large and small, ll: large and large. A breakdown of the performance by object is also shown, with 95% confidence intervals. A total of 80 repetitions were obtained with Subject 2.

